# Importance of eDNA taphonomy and provenance for robust ecological inference: insights from interfacial geochemistry

**DOI:** 10.1101/2023.01.24.525431

**Authors:** K.K. Sand, S. Jelavić, K.H. Kjær, A. Prohaska

**Affiliations:** Section for GeoGenetics, GLOBE Institute, University of Copenhagen, Øster Voldgade 5-7, 1350 Copenhagen, Denmark; Université Grenoble Alpes, Université Savoie Mont Blanc, CNRS, IRD, Université Gustave Eiffel, ISTerre, F-38000 Grenoble, France

## Abstract

**Context for and purpose:** Retrieval of modern and ancient environmental DNA (eDNA) from sediments has revolutionized our ability to reconstruct present and past ecosystems. Little emphasis has been placed, however, on the fundamentals of the DNA-sediment associations and, consequently, our understanding of taphonomy and provenance of eDNA in sediments remains extremely limited. If we are to be able to accurately infer community dynamics across time and space from eDNA data, we need to understand how depositional processes and sedimentary associations of DNA molecules in different settings influence our interpretation.

**Approach and methods:** Here, we introduce interfacial geochemical principles to the field of eDNA and discuss current interpretational biases. We outline a way to increase the scope and resolution of ecological interpretations from eDNA by combining mineralogic composition with experimental adsorption data. We apply distribution coefficients to assess the relationship between the DNA fraction in water columns and DNA fraction sequestered by suspended sediment particles. We further evaluate the challenges with drawing ecological inference using eDNA from sedimentary systems that receive input from different ecosystem types as a consequence of sedimentary processes.

**Main results: We show that:**

- The retention of DNA in aqueous environments depends on the mineralogy of sediment particles and on the number of particles loaded in the water column.
- DNA attached to sediment particles from distal systems can be deposited in proximal systems and skew the interpretation of the proximal sediment samples.
- High particle loading in the water column can deplete suspended DNA and cause inaccurate interpretation of aqueous DNA samples.
- High particle loading in surface sediment pore waters enhances sequestration of DNA from benthic communities relative to that of water column communities, resulting in skewed estimates of species richness and abundance from sedimentary DNA.

We discuss how to integrate taphonomy and provenance knowledge into the reconstruction of modern and past ecosystems, and ecosystem monitoring from eDNA data.

**Conclusions and the wider implications:** Our findings demonstrate that integrating information about eDNA taphonomy and provenance into modern and past ecosystem reconstruction from eDNA data can enhance the scope, resolution and accuracy of our interpretations.

## 1. INTRODUCTION

Environmental DNA (eDNA) holds great promise for improving the spatiotemporal scope of biodiversity research and environmental monitoring. eDNA sampled from air, aerosols, aqueous systems and recent sediments (sedDNA) is increasingly used to obtain information about the composition of present-day communities. Ancient sedimentary DNA (sedaDNA) can provide long-term ecological information within a defined spatial and temporal context. Pairing information derived from sediments about the climatic, physical and chemical depositional environment with sedaDNA-derived community changes across time and space offers a unique opportunity to assess drivers, consequences and tipping points for ecosystems. Despite large developments in the analysis and interpretation of eDNA, however, little emphasis has been placed on DNA taphonomy, i.e. the chemical and sedimentological events and processes that lead to eDNA retention and preservation in sediments, and provenance, i.e. place of origin of DNA or minerals. The field has so far prioritized sampling of sediments from sites that approximate static systems where depositional sources and conditions vary minimally such as closed lake basins or deep ocean floor sediments. However, the importance of understanding eDNA taphonomy and provenance grows with the capability to extract eDNA from ever older and complex deposits and sample sediment sections that increases the likelihood of capturing marked changes that affect the physical and chemical conditions at which eDNA is deposited and stored. If we are to robustly assess changes in ecological communities across time and space using eDNA data, we need to understand how sedimentary processes, their physical and chemical conditions and the mineralogic complexes of DNA molecules influence community reconstruction from eDNA under different environmental forcings.

The eDNA pool is composed of extracellular DNA and intracellular DNA fractions. Here we focus on the extracellular DNA fraction because it is generally more abundant^1^ and more likely to be preserved in the long-term due to the protection it receives upon adsorption to minerals^2^ For instance, Dell’Anno et al. (2005)^1^ estimated that there is ∼0.4 Gt of extracellular DNA associated with the top 10 cm of ocean floor sediments. In general, the majority of DNA in marine benthic ecosystems has been shown to be represented by extracellular DNA rather than living biomass^1,3^ highlighting the importance of surveying this resource.

From interfacial geochemical principles and experiments, we know that the affinity between DNA and minerals changes as a function of aqueous conditions such as pH or salinity.^4–8^ Specifically, adsorption capacity of minerals for DNA is large in seawater because of high salinity that facilitates attractive interaction and lower in freshwater because of relatively stronger repulsive forces. Therefore, the interplay between surface properties of minerals and environmental conditions determines how much of the available dissolved eDNA can be retained and stored in a particular sediment type.^9^ Inherently minerals with a high adsorption capacity for DNA would also be more likely to transport DNA to distant environments. Thus, the interaction between DNA, minerals, solution and depositional conditions will determine which sediments are more likely to carry DNA and influence which DNA-mineral complexes will be preserved in the sedimentary archive.

DNA degradation is directly proportional to the temperature (REFS) so predicting the potential of a sediment to contain eDNA relies on the assessment of the sediment’s ‘thermal history’, i.e., degradation rate of dissolved DNA in a solution^10^ or DNA within a bone.^11^ Consequently, the focus so far has been on relatively young deposits from cold regions because of the higher chance of finding preserved DNA. However, considering interfacial geochemical principles, we hypothesize that the DNA preservation is likely to be influenced by the degree of DNA immobilization through adsorption at a mineral surface. If the DNA is adsorbed through numerous bonds which immobilize the strand, then chemical, biological and thermal degradation is likely to slow down. According to this hypothesis, a thermal DNA degradation model based on kinetics in solution or within a bone is a poor predictor of survival of DNA adsorbed to a mineral. We recently recovered DNA from sediments dated to 2.4 Ma where the thermal age predicted no DNA survival.^12^ Additionally, that study showed that the amount of DNA sediment adsorbs and retains depends on its mineralogic composition. In this communication, the term *preservation* is considered a function of DNA stabilization offered by the substrate (degree of immobilization and binding capacity) as well as resistance to biological, chemical and thermal degradation of the mineral-bound DNA.

Here, we introduce interfacial and geochemical concepts relevant to eDNA taphonomy and provenance, and illustrate how quantitative mineral analysis and DNA adsorption isotherms and dynamics can be applied to assess sources of eDNA and the potential of eDNA preservation in sediments. We use the 2.4 Ma old Kap København sandy deposit as a case study and focus on the following taphonomic processes:

- *Retention of* dissolved DNA at mineral surfaces where we show the importance of *interfacial geochemical processes* (Section 4)
- *DNA partition* between the aqueous solutions and mineral surfaces during and after sedimentation where we stress the importance of mineral loading in a water column for DNA sequestration (Section 5)
- The stability of adsorbed DNA and its relevance for *preservation* where we highlight the co-dependence between mineral surface charge, adsorption and immobilization (Section 6)
- The synergistic influence of eDNA *taphonomy and provenance* on ecological interpretation (Section 7)

We highlight how the information about depositional environments can be exploited to advance the spatial and temporal resolution of ecological inference from eDNA in both modern and past sedimentary deposits as well as in the water column. Collectively, our results demonstrate that by synthesizing mineralogical and sedimentological information with fundamental interfacial geochemical principles in order to characterize eDNA taphonomy and provenance, we can advance the resolution and accuracy of ecological interpretations as well as broaden the range of sedimentary environments where eDNA extraction is feasible.

## 2. OVERVIEW OF INTERFACIAL GEOCHEMICAL CONCEPTS

A major challenge in elucidating the processes that take place in natural systems by using principles of interfacial geochemistry is the immense variability of natural substrates. *E*.*g*., a quartz mineral surface might be partially covered by clay minerals and contain patches of adsorbed humic substances. However, fundamental interfacial reactions take place regardless of the substrate composition. To demonstrate the validity of our argument based on the available literature, we here focus on DNA-mineral interactions. The adsorption capacity of a mineral surface for DNA is related to the mineral surface charge as well as the composition of the mineral surface.^13–15,7,16–19^ The magnitude of the surface charge of most minerals is a function of pH and at pH between 6 -8, silicates are in general negatively charged whereas oxides and hydroxides are positively charged.^20^ A mineral usually has several distinct surfaces each presenting different atoms at the interface, towards the solution. Different surfaces will interact with DNA in different ways, depending on their composition and surface charge. For example, clay minerals, which adsorb an order of magnitude more DNA compared to other minerals,^12^ are composed of layers of silica tetrahedra (T) and layers of magnesium or aluminum octahedra (O) (Fig. 1A)). These layers alternate to form TO, TOT or TOTO structures. Surfaces that make the base of T layers (basal plane) are permanently negatively charged whereas the charge of surfaces at the base of O layers (basal plane) as well as the surfaces at layer edges edges of layers (Te or Oe -edge planes) varies with solution pH (Fig. 1A,B). Thus, negatively charged DNA will interact differently with each of those surfaces.

**Figure 1.**
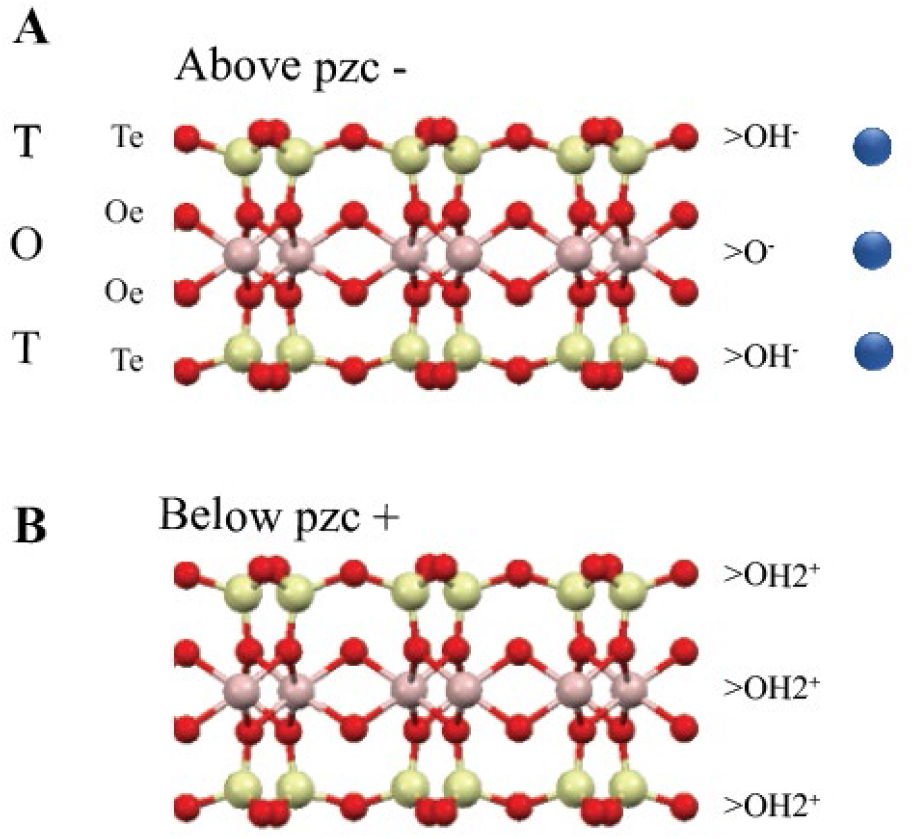
Surface Geochemistry and mineral structure. TOT clay structure showing the speciation of Te and Oe edge sites A) above and B) below the point at which the surface changes its average charge (point of zero charge). Blue spheres: solution cations, red spheres: oxygens, yellow spheres: Si^4+^, pink spheres: Al3^+^. Not to scale.

DNA is adsorbing to mineral surfaces mainly through its phosphate backbone although interactions through the amine groups on the bases are also shown to occur.^18,21–23^ Binding can occur electrostatically, through hydrogen bonds or van der Wahl bonds^23,24^. Among those, the electrostatic bonds are in general the strongest. Here a negatively charged DNA molecule binds directly to a positive mineral surface, *i*.*e*. to positively charged clay edges or O layers. Above pH 5 where the phosphates moieties of DNA are overall negatively charged, adsorption of DNA at negatively charged mineral surfaces (negative edges or tetrahedral basal planes) rely on a screening of charges by cations. Depending on the ionic potential of the cations (charge/radius), the adsorption capacity varies greatly and is enhanced by the presence of polyvalent cations such as Ca^2+^, Mg^2+^ or Al^3+^ which require lower concentrations to screen the repulsive forces than monovalent cations such as Na^+^. ^23,24^

The consequences of the influence of solution chemistry on the DNA-mineral complexes is important for eDNA interpretations in several aspects. First, the adsorption capacity of minerals for DNA was found to be sensitive to the solution (types of ions,^25,26^ salinity and pH^4–8^) and to vary among minerals.^13–15,7,16–19^. The affinity of a mineral for DNA can be described by adsorption isotherms that provide a relationship between the adsorbed DNA (adsorption capacity) and DNA that was left in solution (equilibrium concentration, *c*_*eq*_) once the equilibrium in adsorption was reached. Reported concentrations of dissolved DNA in seawater typically range from 0.03 to 88 μg per L, and decrease as a function of distance from the shore and the depth of the water column.^27–31^ Removal of suspended DNA through mineral adsorption is particularly relevant in environments where the concentration of mineral particles in the water column (the suspension loading) is large or it varies considerably as a function of depositional characteristics, *e*.*g*., in coastal regions where the terrestrial particle influx. Yet the sequestration of suspended DNA to minerals has not been given much attention to date.^32–34^

## 4. CASE STUDY: The Kap København Formation

### Mineralogy and adsorption characteristics

The Kap København formation is a 2.4 Ma marine sandy deposit that contains the oldest DNA recovered to date.^12^ The sediment profile can be divided in two depositional environments with similar overall mineral composition but of varying ratios. Excluding the amorphous fraction (both organic and inorganic), the formation contains on average 69 wt% of quartz and 7.3 wt% of clay minerals: the rest of the major minerals are feldspars, pyroxene and amphiboles (Fig. 2) and Sup.Table S1). The isotherms describing DNA adsorption to the individual minerals in seawater are shown in Figure 2. The clay minerals have a much higher DNA adsorption capacity compared to quartz, feldspars, pyroxene and amphiboles. The adsorption capacity of a mineral was found to be roughly inversely proportional to the recovery yield: more than 40% of the adsorbed DNA could be recovered from quartz and feldspars, but only 5% could be recovered from smectite, the mineral with the largest adsorption capacity for DNA.

**Figure 2.**
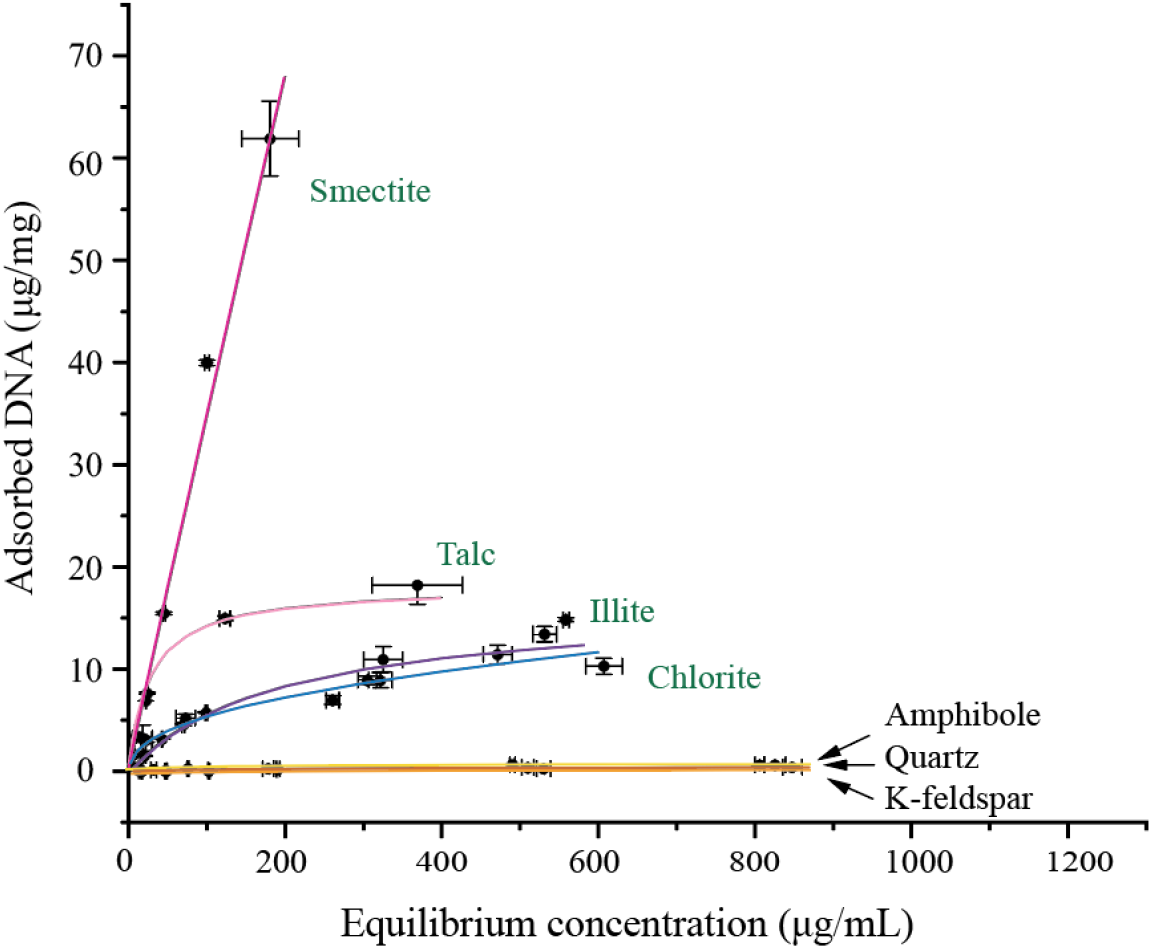
Adsorption capacity of major minerals from Kap København formation.^12^ The adsorbed DNA is shown as a function of the DNA equilibrium concentration, i.e. the concentration of DNA in solution once the equilibrium was reached. The uncertainty is expressed as a standard deviation of three measurements. The colored lines represent a fit to the Freundlich adsorption model (chlorite and smectite) or the Langmuir adsorption model. Names of clay minerals are green.

### Assessing the DNA adsorption and storage capacity of Kap København deposit

We translate the adsorption capacities and the quantitative mineral composition of a Kap København sediments into the following indicators:

1. Normalized adsorption capacity, which describes how much extracellular DNA a mineral in a deposit can retain relative to the other minerals a) assuming an equal concentration of minerals and b) normalizing the adsorption capacity to 1 (equation in Supporting Information section 2). The normalized adsorption capacity show which mineral retains most DNA at a given DNA equilibrium concentration,and is information that can guide the choice of sampling strategies and extraction protocols, as well as interpretation of eDNA data based on the mineral’s provenance.
2. DNA storage capacity, which describes how much extracellular DNA each mineral in a specific sediment can store considering the actual weight percent of the mineral in the deposit (equation in Supporting Information section 2). Knowledge of which mineral the majority of the DNA is associated with can help guide the choice of extraction protocols as well as sampling, subsampling and interpretations. Combined with information on the depositional environment, the storage capacity can provide an approximation of which species were native to the depositional environment, and which could be considered introduced.
3. Extraction yield, which describes how much of the adsorbed DNA can be extracted using the phenol -chloroform method.^35^

The values of these three indicators are visualized in Fig. 3. The composition displayed in Fig. 3A represents the average mineral composition of Kap København formation. Merging the mineralogic composition with adsorption capacity, we get an estimate of how much DNA each mineral in the deposit has the capacity to store. In Fig 3B-D, we consider two distinct extracellular DNA concentrations to represent a) a DNA-poor environment in the Arctic Beauford Sea (*c*_*eq*_ = 0.0049 ug/mL)^36^ and b) an environment with a high DNA concentration (*c*_*eq*_ = 200 ug/mL), to show the influence DNA concentration has on the relative adsorption capacities. The considered *c*_*eqs*_ are chosen to illustrate the influence of *c*_*eq*_ on the normalized adsorption and storage capacities of a deposit. We show that regardless of the *c*_*eq*_ chosen, the clay minerals adsorb at least 98.6% of the total adsorbed DNA (minerals >0.6 wt% were included in adsorption experiments) (Fig. 3B and Table S2.2). Only 0.2 - 1.4% of the total DNA concentration is associated with the non-clay minerals, regardless of the large range of considered *c*_*eq*_. Differentiating between the clay minerals in the Kap København formation (bar diagrams imposed on the “treemaps” in B-D) the clay mineral that are able to adsorb and store the most DNA depends on *c*_*eq*_. At a low *c*_*eq*_, chlorite has the highest normalized adsorption (and storage capacity) where 70% of total adsorbed DNA was adsorbed to chlorite, whereas smectite-adsorbed DNA accounts only for 2.6% (Fig 2C). However, at higher *c*_*eq*_, most DNA can be stored by smectite. Considering the amount of DNA we were able to extract using the phenol-chloroform method, the DNA adsorbed to clay minerals became less represented in the extract, especially at higher *c*_*eq*_. Only about 10% of the total storage capacity was extractable from the clay minerals. With the low recovery of the DNA adsorbed to clay minerals, the amount of DNA adsorbed to quartz, feldspar, pyroxenes and amphiboles only becomes significant at high C_eq_ (Fig 2D).

**Figure 3A-D.**
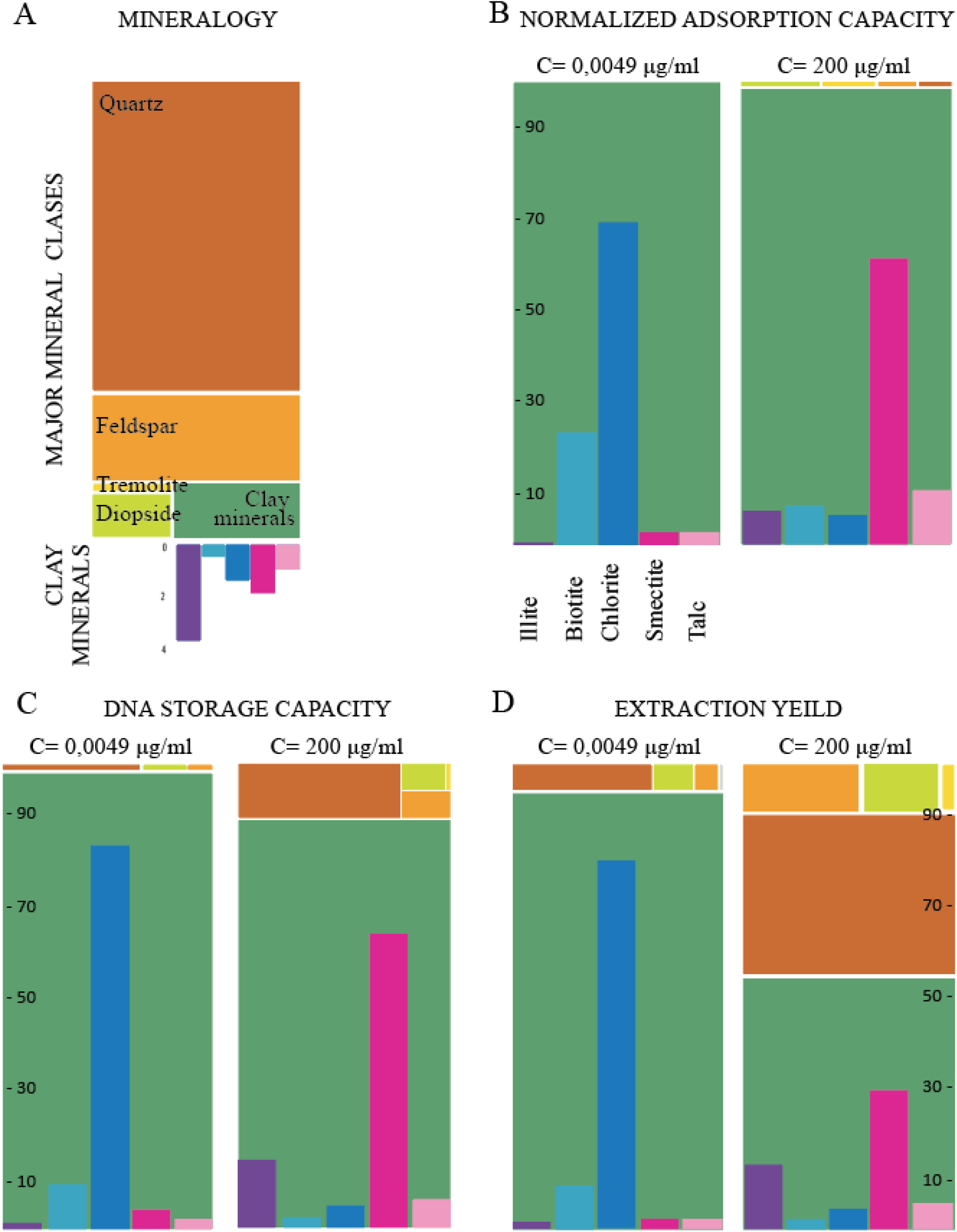
Indicators for the DNA adsorption and storage capacity. A) Mineral composition in wt% of the Kap København deposit (only minerals >0.6 wt% taken into account, excluding amorphous components). The bar diagram below the clay mineral block in the tree diagram represents concentration of individual clay minerals in the deposit (mineral names in 2B, applies to A - D). Tree diagrams in % of DNA, in B) normalized adsorption capacity, and C) storage capacity, and D) extraction yield for a low and a high equilibrium concentration of DNA (*c*_*eq*_). The concentration of each clay mineral in % is shown as bar diagrams superimposed on the clay mineral block.

## 5. DNA SEQUESTRATION DURING SEDIMENTATION

### Partitioning of environmental DNA between the aqueous environment and mineral surfaces

In principle, the capacity of minerals to sequester DNA from seawater depends on the mineral:water ratio and the mineral composition of suspended particles in the water column. The mineral:water ratio in the water column is called the particle loading and can be envisaged as a mass concentration of particles in a kilogram of water. Loading varies between different marine settings; coastal regions typically have a high particle loading because of high riverine input, and deep-water environments have a relatively lower particle loading with an increased aeolian input. Such varied loading distribution within a depositional environment combined with different particle provenance likely impacts species composition inferred from eDNA data. In the next few lines, we look into differences in DNA sequestration between coastal and deep-water regions, proxied by varied particle loading, and between particles of different provenance, proxied by different mineral compositions.

First, we will consider the DNA that is left in the water column after the adsorption has reached equilibrium (equilibrium DNA, *c*_*eq*_) because it is an intuitive measure of DNA fate in seawater. We can express *c*_*eq*_ using a distribution coefficient:

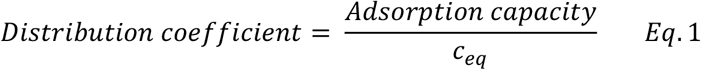

where adsorption capacity represents DNA adsorbed to a mineral surface. For this calculation, we assume that a) all particles in a seawater column are made of a single mineral species, b) extracellular DNA is double stranded and c) the influence of other organic compounds such as proteins and lipids on DNA adsorption is negligible. Inserting the distribution coefficient into the equation for adsorption capacity (Supporting Information section 3), we can calculate the fraction of DNA in seawater as a function of a loading:

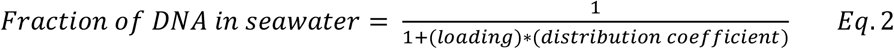

The fractions of DNA in seawater with different particle loadings and for two different *c*_*eq*_ are shown in Table 1. For silicates such as quartz, feldspar, amphibole and pyroxene that have small adsorption capacity for DNA, a vast majority of DNA is left in the water column after adsorption has reached equilibrium irrespective of the *c*_*eq*_ and the particle loading. The behavior is similar for clay minerals but some such as talc and smectite are capable of removing up to 10 - 15% of DNA from the water column at high particle loadings. An exception to this is chlorite that can remove the majority of extracellular DNA from seawater with high particle loading (for a *c*_*eq*_ = 0.0049 μgml^-1^).

**Table 1.** DNA fraction left in seawater after adsorption in the water column for three different particle loadings and two different DNA equilibrium concentrations, *c*_*eq*_. Values rounded to no decimal places.

|  | Fraction of DNA in seawater (%) |  |  |  |  |  |
| --- | --- | --- | --- | --- | --- | --- |
| | $c_{eq}(\text{DNA}) = 0.0049 \mu\text{gml}^{-1}$ | | | $c_{eq}(\text{DNA}) = 200 \mu\text{gml}^{-1}$ | | |
|  | Far from coast<sup>37</sup><br>(North Sea) | Coastal region<br>(North Sea) | High-load sea<sup>38</sup><br>(Yellow + East China Sea) | Far from coast<br>(North Sea) | Coastal region<br>(North Sea) | High-load sea (Yellow + East China Sea) |
| Quartz | 100 | 100 | 100 | 100 | 100 | 100 |
| K-Feldspar | 100 | 100 | 100 | 100 | 100 | 100 |
| Amphibole | 100 | 100 | 100 | 100 | 100 | 100 |
| Pyroxene | 100 | 100 | 99 | 100 | 100 | 100 |
| Talc | 100 | 94 | 84 | 100 | 99 | 98 |
| Illite | 100 | 99 | 98 | 100 | 100 | 99 |
| Kaolinite | 100 | 97 | 92 | 100 | 100 | 99 |
| Chlorite | 97 | 40 | 18 | 100 | 100 | 99 |
| Smectite | 100 | 96 | 88 | 100 | 97 | 91 |

For sediments at the ocean floor, the mass concentration of particles is very high compared to the water column, to the extent where the term particle loading is not appropriate anymore, so we define porosity as a measure of mineral:water ratio. The porosity is a fraction of pore spaces (voids) in a volume of sediment so the fraction of DNA in sedimentary pore waters can be calculated by relating the porosity to loading:

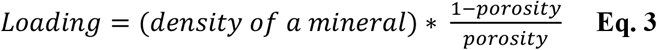

and recasting eq 3 into eq 2.

Table 2 shows the DNA fraction in pore waters in sediments such as beach sand or deep marine mud where the porosity ranges from 0.3 – 0.9.^39^ Unlike in the water column, the vast majority of DNA is removed from pore waters after the adsorption equilibrium is reached even for low adsorbing minerals such as quartz, feldspars, amphiboles and pyroxenes, except in high porosity sediments that can still leave a considerable fraction of DNA in the solution. Table 2 shows that clay minerals are more efficient at removing DNA from pore waters than non-clay minerals, irrespective of the sediment porosity.

**Table 2:** Fraction of DNA left in a pore water for two different porosities and two different DNA equilibrium concentrations, *c*_*eq*_.

|  | Fraction of DNA in pore water (%) |  |  |  |
| --- | --- | --- | --- | --- |
| | $c_{eq}(\text{DNA}) = 0.0049 \mu\text{gml}^{-1}$ | | $c_{eq}(\text{DNA}) = 200 \mu\text{gml}^{-1}$ | |
|  | Low porosity* | High Porosity^ | Low porosity | High porosity |
| Quartz | 5 | 53 | 10 | 71 |
| K-Feldspar | 7 | 63 | 16 | 80 |
| Amphibole | 3 | 36 | 7 | 63 |
| Pyroxene | 1 | 16 | 5 | 52 |
| Talc | 0 | 1 | 0 | 5 |
| Illite | 0 | 4 | 0 | 8 |
| Kaolinite | 0 | 1 | 1 | 14 |
| Chlorite | 0 | 0 | 0 | 9 |
| Smectite | 0 | 1 | 0 | 1 |

\* Porosity of a beach sand taken at the low limit as 0.3
^ Porosity of a deep marine mud taken at the high limit as 0.9

Despite the simplistic assumptions applied in the calculation of the distribution coefficient, Table 1 and 2 demonstrate the importance of minerals in DNA sequestration from seawater in different marine environments. For example, a beach sand with a low clay mineral content would essentially deplete the pore water from DNA. On the other hand, a river plume adsorbs only a tiny fraction of dissolved extracellular DNA from seawater even if all the particles adsorb large amounts of DNA such as clay minerals. The only exception to this is a river plume made of chlorite particles, which can theoretically remove more than 50% of DNA from the water column. The fact that DNA from pore waters is depleted relative to the DNA from the water column suggests that the benthic species will be overrepresented in the DNA extract compared to pelagic species, hence the relative proportions of species abundance will be skewed.

## 6. STABILITY OF MINERAL ADSORBED DNA

When evaluating the preservation potential of ancient eDNA in a particular deposit, it is useful to distinguish between the interfacial, and the chemo-biological component of preservation. The interfacial component is the capability of a mineral to retain DNA. The chemo-biological component is the resistance of DNA to degradation which is manifested as the resistance to UV, hydrolysis or enzymatic degradation. In the following, we focus on the interfacial component and use AFM to image the DNA during adsorption at chlorite surfaces.

Chlorites are a group of clay minerals characterized by high negative charge of the TOT layers that is neutralized by an additional positively charged octahedral layer, not present in other clay mineral groups. The unit layer of chlorites is thus called a TOTO layer. The negatively charged TOT layers (Fig. 1A) are structurally similar to mica and are 1.0 nm thick. The additional positively charged octahedral layer is ∼450 pm thick, and structurally and compositionally similar to brucite (Mg(OH)_2_). The structure and the two types of surface charging make chlorite a reasonable model-system to illustrate how the adsorption of DNA at mineral surfaces changes as a function of mineral surface composition. Figure 4A shows irregular bands of mica-like and brucite-like sheets at the chlorite surface.^40,41^ The irregular bands occur because of the difference in topography, where brighter colors designate topographically higher parts of the AFM image. The height profile in Fig. 4B shows that the brighter areas (terraces) in Fig 4A are ∼450 nm tall and are hence the brucite-like sheets. The darker areas are then the mica-like sheets. In artificial seawater (ASW) at pH=7, plasmid DNA mainly adsorbs to the positively charged brucite-like sheets and the DNA adopts two forms: smaller blobs scattered around the brucite-like terraces (Fig. 4 C, D, E) and the fully stretched-out form that adsorbs at the terrace step edges (Fig. 4 F). Occasionally the plasmids are found anchored to the blobs on the terraces (arrow in Fig 4D, E). Snapshots from near-video-rate imaging in Fig. 4D and 4E show that its uncoiled strand is mobile and associates with different blobs. The mobility of the strand during the image acquisition shows that it is not as strongly bound by the bulk of the brucite-like surface compared to the DNA that is immobilized at the step edges during image acquisition (Fig. 4F). A stronger binding at the step edges is a result of higher charge density compared to the terraces, as observed at calcite step edges as well (Freeman et al., 2020). The strands bridging between two brucite-like step edges across the mica-like valley are immobile at the step edge but mobile at the mica-like surface (Fig. 4G-I). The absence of DNA adsorbed exclusively to the mica-like sheets is likely an artifact of the imaging approach because the binding of the double stranded DNA to the negatively charged mica is likely weak enough that, during the imaging, the lateral tip forces perturb the adsorption equilibrium and displace the DNA (Supporting information section 4).

**Figure 4.**
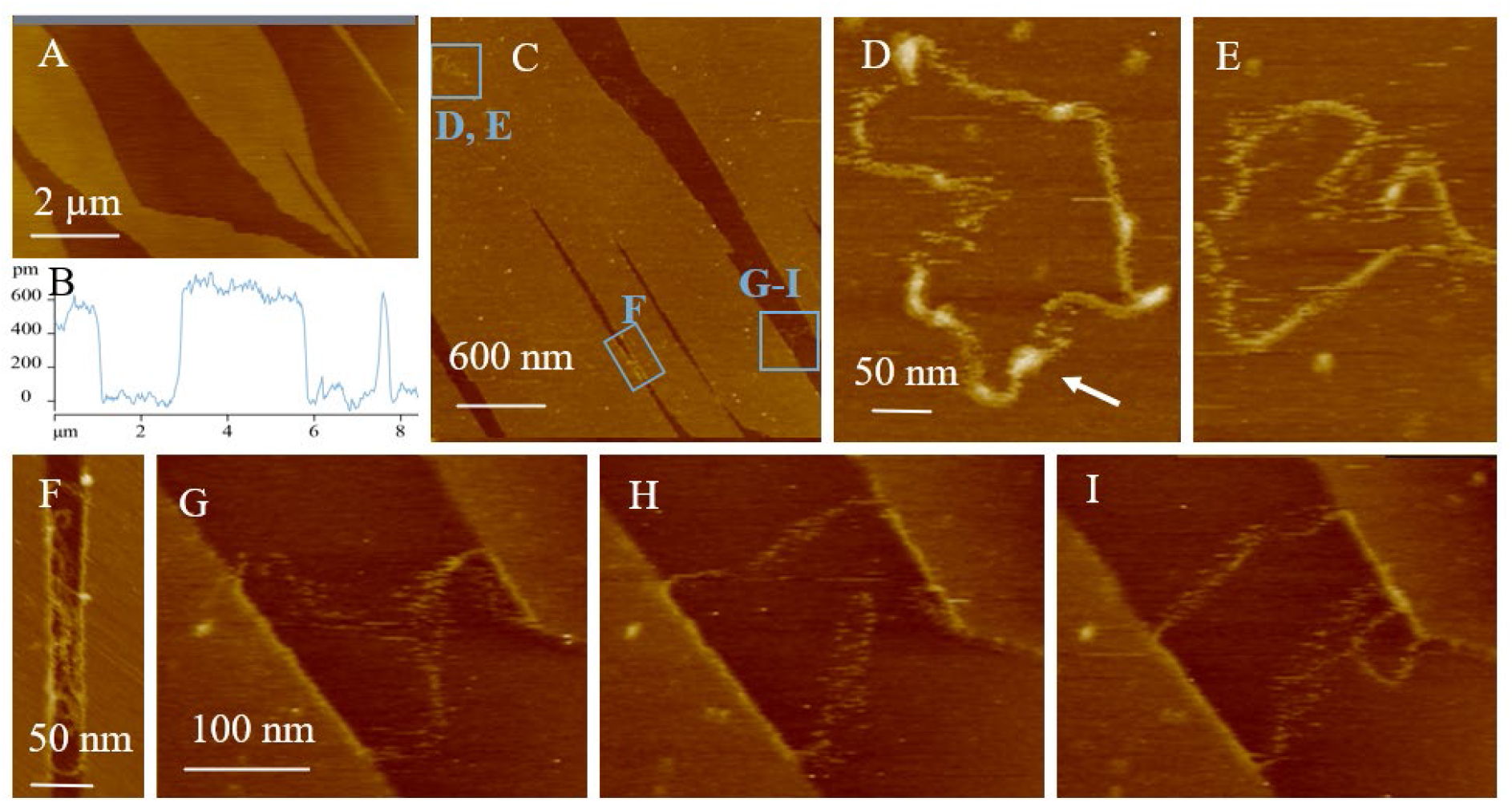
Atomic force microscopy images of chlorite and plasmid DNA interactions. A) Chlorite surface imaged in air. The height profile (B) shows positively charged octahedral brucite-like sheets (450 nm thick) as bright terraces and the tetrahedral mica-like sheets as darker valleys. Adding plasmid DNA in artificial seawater (ASW) shows distinct behavior of the plasmids depending on the local composition: either brucite- or mica-like (C). D-I) show the inserts indicated with blue in C. D) and E) show the same plasmid molecule interacting with the octahedral layers at two different time steps 40 s apart). Occasionally the uncoiled plasmid associates with the coiled plasmid (arrow). The plasmid is relatively stronger bound to the brucite-like step edges compared to strand fractions that bridge across the mica-like valleys, which are highly flexible, thus bound relatively weakly (F, G-I).

Fig. 4 demonstrates that the increase in surface charge density improves the immobilization of DNA. The immobilization can be explained by relatively stronger binding and a larger number of bonds between the DNA and mineral surfaces. Consequently only a small portion of DNA strand is likely to be available for hydrolysis. Further, the configuration of the DNA is modified by adsorption which is thought to prevent enzymatic recognition^42,43^ and subsequent degradation. The implications are that DNA adsorbed to negatively charged minerals (silicates and clay mineral basal planes) are likely to degrade faster compared to DNA adsorbed to charge dense areas carrying a positive charge such as terrace steps and edge sites. As an example, it is well established that cations with a high ionic charge density such as Ni^2+^ are able to immobilize DNA on a negatively charged mica surface and change the DNA conformation, probably because the charge dense Ni^2+^ can provide several strong adsorption sites for the DNA.^44^ The importance of cations for adsorption and, ultimately, immobilization of DNA to negatively charged mineral surfaces render the preservation of such complexes more sensitive to solution composition than DNA directly or strongly bound to a positively charged surface.

## 7. INFLUENCE OF TAPHONOMY AND PROVENANCE FOR INTERPRETATIONS OF BIODIVERSITY AND PALAEOECOLOGY

Knowledge on the parameters controlling DNA preservation on minerals and on the DNA extractability can facilitate the interpretation of species richness and abundance from both sedimentary and water eDNA. The adsorption of DNA on minerals described in Kap Købenahvn formation shows that clay minerals have higher DNA adsorption capacity compared to non-clay minerals.^12^ Following the principles of interfacial geochemistry, DNA adsorption to positive sites (at neutral pH) such as clay mineral edges is direct, thus will be less influenced by salinity and cationic composition than DNA adsorption to non-clay minerals. Because of generally high surface area of clay minerals and positive charged edges, they are more likely to contain DNA from distal environments than other silicates. The concentration of DNA transported from distal environments might be considerable if the minerals have been reworked from a previously deposited site.

As a consequence the taphonomic processes, inference of species abundance from sedaDNA will be influenced in several ways: a) the origin of the DNA, *i*.*e*., whether it originated from the environment from which it was sampled in or from another environment, b) the timing of mineral-DNA complex formation, *i*.*e*. pre-, syn- or post- sedimentation, c) the loading of suspended minerals in the water column (Table 1), d) the porosity of the sediment (Table 2), and e) the resistance to degradation of the adsorbed DNA, *i*.*e*. weakly bound DNA is less likely to be preserved than strongly bound DNA. In Fig. 5 we consider these factors and illustrate how eDNA-based species abundance obtained from marine sediments may not reflect the actual local abundance (Fig. 5). Our results further highlight that when using eDNA data to reconstruct biodiversity changes across time and space, comparison will be most robust among sites with similar geological and taphonomic conditions. We illustrate our hypothesis through the comparison of two models - a coastal and a deep marine environment, where we, in the latter, assume that the concentration of dissolved extracellular DNA is representative of the living biomass, *i*.*e*. relatively low.

**Figure 5.**
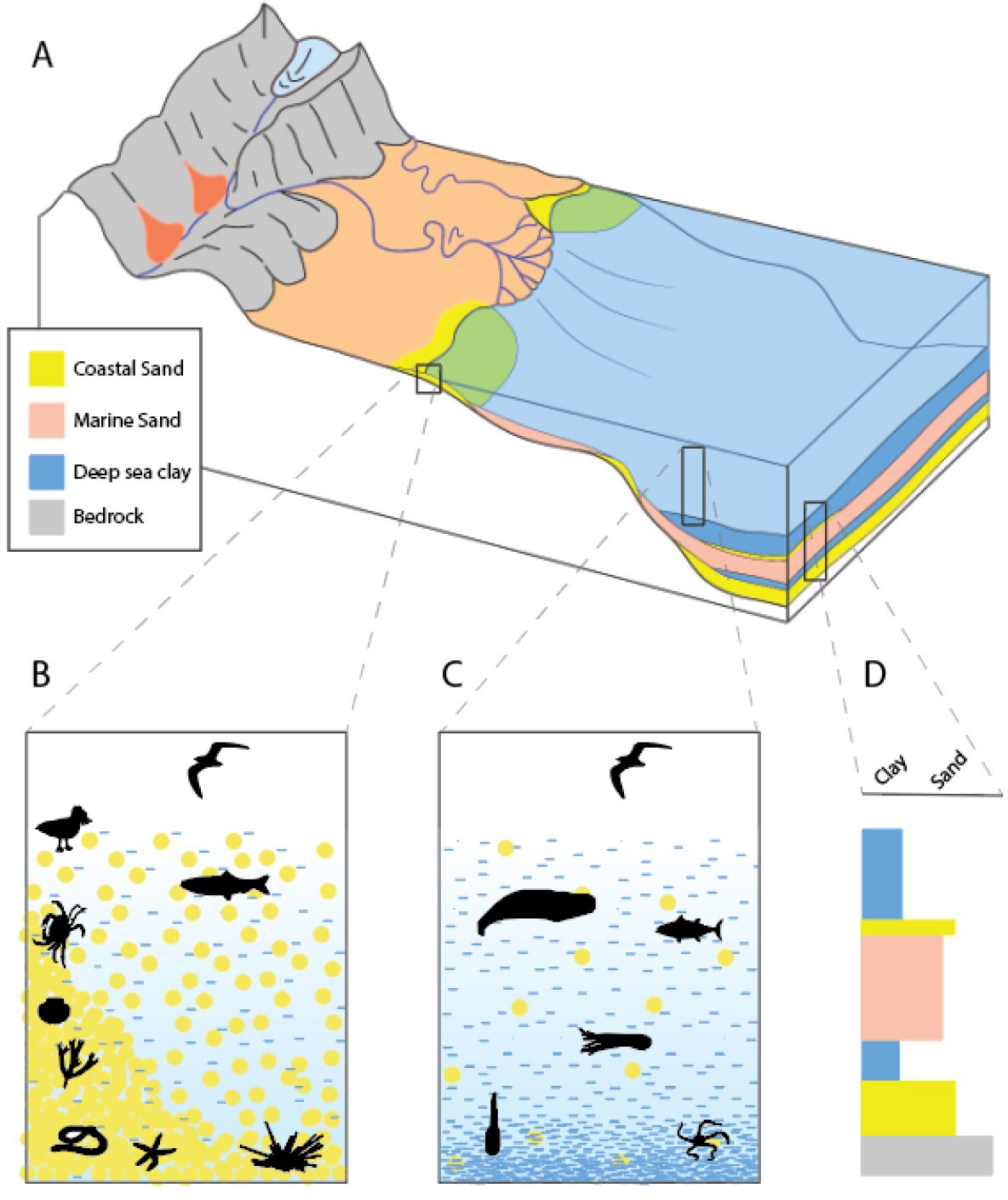
A) Schematics of depositional settings showing the differences between a coastal site (B) and a deep marine site (C). At the coastal site there is a rich benthic environment, and the sediment is mainly composed of quartz (yellow) while most clay minerals (blue lines) are present in the water column. At the deep marine site, the sediment is enriched in clay minerals with relatively little quartz and the sedimentation rate is lower than at the coastal site. D) shows a section of the stratigraphy under the modern deep sea clay depositional setting. The section illustrates that distinct depositional environments are superimposed on each other and highlights that the difference in identified taxa between successive layers is a mixed signal of the depositional environment and derived changes in biodiversity.

Let us assume that the coastal and deep marine sediment displayed in Fig. 5 contains quartz and clay minerals. Clay minerals are small in size (on average <2 μm in diameter) so they will be enriched in deep marine sediments because they can be carried further away by the currents, unlike quartz particles that are larger on average and will settle closer to the shore. However, it is important to recognise that clay minerals and quartz coexist in a suspension while traveling outbound in a high-energy current such as those present in rivers. The coastal and deep marine environment host different ecological assemblages inherent to the present ecological niches (5B vs. 5C).

At the coastal site (Fig. 5B), clay minerals, quartz and other non-clay silicates coexist in the water column, however quartz is predominately deposited. The environment is of too high energy for the clay minerals to settle and they are to a great extent transported to deep waters where they comprise a clay deposit (5C) with a low sedimentation rate. For particle loadings representative of a coastal site (Table 1), clay minerals adsorb only minor concentrations of dissolved DNA (except chlorite that can adsorb up to 60%) at *c*_*eq*_ = 0.0049 μgml^-1^ while quartz and other non-clay minerals adsorb only traces. However, because of the high mineral:water ratio within a sediment, the quartz in the sand adsorbs most of the DNA from the pore waters. Consequently, the DNA adsorbed by quartz likely consists of a smaller fraction of DNA from organisms in the water column from the catchment and the continental shelf, and a larger fraction of DNA from the benthic organisms.

Following the clay mineral deposition at deeper waters (Fig. 5C) where the energy regime is low, the low particle loading in the water column only allows trace amounts of DNA to adsorb (Fig. X2). DNA adsorbed to mineral particles from the water column is more likely to originate from the coastal area because of the higher mineral loading compared to the pelagic zone. In addition, the input of DNA from pelagic species to the reservoir is lower relative to species from coastal areas because of generally fewer inhabitants in the deeper water column. After sedimentation, the clay minerals will essentially deplete the pore water from dissolved DNA (Table 2). Collectively, the species identified from the deep marine sediments will be a mixture of species found in catchment and in the coastal zone, within the water column and within the sediment with an overrepresentation of benthic species.

The inherent differences in the sedimentation processes of the two sampling environments has important implications for inferring changes in biodiversity over time and space. Assuming a similar preservation potential at two depositional sites, the coastal site will provide a more local biodiversity record, whereas the DNA retrieved from the deep marine site will give a more regional signal. The eDNA records of both the coastal and the deep marine site will have overrepresentation of benthic taxa, which may be misinterpreted as ecological dominance rather than sampling and geochemical bias. The eDNA record from either site can, in principle, provide realistic estimates of relative species abundances within benthic communities. Abundance as well as dissimilarity measures such as turnover and nestedness can be compared to other geologically similar sites, *i*.*e*. between similar layers of deep-marine strata in a sedimentary column (Fig 4d). In contrast, comparing ecological communities in successive layers of different sediment types might give inaccurate measures of biodiversity changes.

Benthic community sampling is already an established practice of environmental eDNA monitoring. Considering that extracellular DNA from benthic communities are well sampled by the local sediments, benthic diversity assemblages may provide a solid approach for tracking changes in biodiversity across time and space in both “static” and dynamic settings. It is vital though that the sediment provenance and history is considered to avoid misidentifying transported benthic DNA as local DNA. In a fluvial environment (Fig 5 III) sediments will have been redeposited numerous times and thus samples of a fluvial-run-off sediment will carry a signature of fluvial benthic taxa.

In terms of biodiversity monitoring in modern settings using extracellular DNA, the mineral loading in the water column as well as the mineral composition will influence the extractable extracellular DNA, either adsorbed or dissolved. The influence of particle loading and mineral composition on DNA retention likely explains some inconsistencies in community composition reported in the literature.^45^ For instance, aqueous sampling for fish DNA has been shown to yield a higher diversity compared to fish DNA from sediment.^45^ Likewise, contradictory results about the turnover rate of extracellular DNA originating from the organisms living in the water column and sampled from the sediment below^45–48^ could be at least in part explained by different sediment input and concomitant DNA retention from lagoons, coastal regions and open sea.

## 8 CONCLUSIONS AND OUTLOOK

Our results demonstrate that eDNA taphonomy and provenance is heavily shaped by the interactions between the extracellular DNA and the mineral surface, and pre-, syn- and post- sedimentological processes. Furthermore, our findings highlight the enormous potential of mineralogic analysis, interfacial geochemical principles, and consideration of sedimentological processes for advancing the understanding of eDNA taphonomy and provenance.

We recommend the use of mineralogical and sedimentological analyses to:

- unlock promising new reservoirs of ancient eDNA,
- develop sound eDNA sampling strategies,
- identify optimal DNA extraction procedures,
- improve ecological interpretation from eDNA data.

The knowledge of sediment retention capacities and mineral loading, and its integration with the information about the relationship between distal and proximal depositional environments should go a long way in improving the resolution and scope of both present-day biodiversity assessments and ancient ecosystem reconstructions.

## ACKNOWLEDGEMENTS

This work was supported by a research grant (00025352) from VILLUM FONDEN.” SJ was partly funded by French Government through MOPGA Postdoctal Programme (ref. 3-5402234721) and LABEX

## CONFLICT OF INTEREST STATEMENT

The authors declare no conflict of interests.

## AUTHOR CONTRIBUTIONS

KKS developed and conceptualized the ideas, made the AFM experiments, developed the concepts of adsorption and storage capacity, placed the data in relevance of taphonomy and provenance for ecological interpretations and wrote the original manuscript. SJ conceptualized the ideas, developed the relationship between the mineralogy DNA adsorption and sequestration, contributed to implementing it in the discussion about its ecological framework. AP provided the ecological context to the study and helped frame its broader relevance. KK provided context for depositional environments and Figure 5. All authors were involved in manuscript writing.

## MATERIALS AND METHODS

### DNA-mineral adsorption dynamics

We used atomic force microscopy to study the DNA chlorite interaction at the nanoscale. The chlorite was freshly cleaved using tape. The salts for the artificial seawater (ASW) solution^49^ were reagent grade and purchased from Sigma Aldric and used without purification. The double stranded plasmid DNA (pUC19) was purchased from Integrated DNA Technologies IDT and annealed prior to use. All solutions were prepared using molecular grade biology water. For DNA adsorption to the chlorite in air, we added a 10 μL droplet with pUC19 at a concentration of 0,5 ng/μL to the surface for 1 min. The droplet was rinsed with 400 μL double filtered Molecular grade water to remove salts. The mineral was dried with a soft blow of N_2_(g) for 30 seconds. For scanning in air and liquids, we used the non-contact mode on a Cypher and a Cypher VRS from Oxford Instruments. Both were running in tapping mode. Aluminum coated AC240TS silicon tips from Olympus with nominal spring constants between 0.6 and 3.5 N/m were used for imaging in air and BL-AC40DS from Asylum Research, Oxford Instruments were used for imaging in liquid. We used a spring constant of 107 pN/nm and a resonance frequency of 1500 kHz. The spring constants were determined using the Sader method (Sader et al. 2012). For both instruments, the force between the tip and the sample was varied to minimize the force exerted by the tip on the surface.

## Notes

### Competing Interest Statement

The authors have declared no competing interest.

