## Supporting Information for "Importance of eDNA taphonomy and provenance for robust ecological inference: insights from interfacial geochemistry"

#### Supporting info section 1. XRD

Table S1. Average mineralogical composition, adsorption data and extraction recovery

| Unit | Average (wt%) | Equivalent applied in adsorption study | *$q_{max}$ Langmuir* | *$K_F$ Freundlich* | *Extraction Recovery (%)* |
|---|---|---|---|---|---|
| Quartz | 68.57 | Quartz | 0.46 | / | 43 ± 32 (7) |
| K-feldspar | 10.61 | Orthoclase | 0.26 | / | 43 ± 28 (3) |
| Plagioclase | 8.31 | / | / | / | |
| Hornblende | 0.80 | Tremolite | 0.6 | / | 43 ± 17 (3) |
| Diopside | 4.00 | Diopside | 0.73 | / | 36 ± 10 (3) |
| Rutile | 0.20 | / | / | / | / |
| Magnetite | 0.10 | / | / | / | / |
| Halite | 0.10 | / | / | / | / |
| Pyrite | | / | / | / | / |
| Gypsum | | / | / | / | / |
| TOTAL non-clays | 92.69 | / | / | / | / |
| Dioctahedral mica | 3.31 | Illite | 16.47 | / | 10 ± 2 (4) |
| Trioctahedral mica | 0.39 | / | / | / | |
| Trioctahedral chlorite | 1.20 | Chlorite | / | 0.73 | 10 ± 1 (3) |
| Trioctahedral smectite | 1.61 | Smectite | / | 0.40 | 4 ± 1 (4) |
| Talc | 0.80 | Talc | 15.84 | / | 10 ± 2 (4) |
| TOTAL clays | 7.31 | | | | |

Minerals with an average of 0.6 wt% or more were included in the adsorption study. For a full list of adsorption parameters see Kjær et al. 2022. The high uncertainties on extractions recovery from non-clay minerals relates to their low adsorption capacities for DNA.

### Supporting info section 2: Retention

For assessing DNA retention in sedimentary deposits, it is practical to normalize the mass of adsorbed DNA to the mass of a mineral component since the weight % of a mineral in a rock or sediment is easier to determine than its specific surface area. Focusing on a specific deposit we can normalize the adsorption capacities of the minerals present to 100 and obtain a measure that can be related to the wt% of the quantified minerals (normalized adsorption capacity of each of the minerals in a deposit). We can then use the relative wt% of the minerals present and the normalized adoption capacity for each mineral to access how much DNA each unit of mass of the different minerals can store at a specific DNA $C_{eq}$. Such measure of storage capacity of the individual minerals in a sediment indicate which minerals are the major and minor DNA carriers in a particular deposit. Knowing which mineral is associated with the majority of the DNA, we can further assess if an extraction protocol optimized for, *e.g.*, clay minerals could be beneficial.

Calculations of adsorption and storage capacities,

We here provide equations for calculating the adsorption capacities, the normalized capacities and storage capacities for each mineral in the Kap Copenhagen deposit. The equations are applicable when the mineralogic composition of a sediment is known and valid under the assumption that the mineral surfaces are clean, *i.e.,* no previous DNA or other organic compounds are adsorbed.

The adsorption capacities of the studied minerals are defined by a fitted isotherm (Ref to kap K) and can for each mineral be expressed as:

Adsorption capacity (Quartz) = $\frac{0.003c_{eq}}{1+0.006c_{eq}}$

Adsorption capacity (K-feldspar) = $\frac{0.002c_{eq}}{1+0.007c_{eq}}$

Adsorption capacity (Amphibole) = $\frac{0.006c_{eq}}{1+0.010c_{eq}}$

Adsorption capacity (Pyroxene) = $\frac{0.017c_{eq}}{1+0.023c_{eq}}$

Adsorption capacity (Talc) = $\frac{0.619c_{eq}}{1+0.040c_{eq}}$

Adsorption capacity (Illite) = $\frac{0.082c_{eq}}{1+0.005c_{eq}}$

Adsorption capacity (Kaolinite) = $\frac{0.301c_{eq}}{1+0.070c_{eq}}$

Adsorption capacity (Chlorite) = $0.73c_{eq}^{0.43}$

Adsorption capacity (Smectite) = $0.40c_{eq}^{0.97}$

Following the adsorption capacities, the mass of adsorbed DNA per mass of a mineral ($\mu g mg^{-1}$) equilibrated in seawater can be calculated. TableX show the adsorbed DNA concentrations ($\mu g mg^{-1}$) for two different $C_{eq}$.

**TableS2.1**: Adsorption capacities at two different DNA equilibrium concentrations ($C_{eq}$) based on adsorption isotherms measured in ASW.

| | Adsorption capacity (**µgmg$^{-1}$) at** | |
|---|---|---|
| | $C_{eq}$ **= 0.0049 µgml$^{-1}$** | $C_{eq}$ **= 200 µgml$^{-1}$** |
| Quartz | 1.5 x $10^{-5}$ | 0.273 |
| K-Feldspar | 1.0 x $10^{-5}$ | 0.167 |
| Amphibole | 2.9 x $10^{-5}$ | 0.400 |
| Pyroxene | 8.3 x $10^{-5}$ | 0.607 |
| Talc | 3.0 x $10^{-3}$ | 13.756 |
| Illite | 4.0 x $10^{-4}$ | 8.200 |
| Kaolinite | 1.5 x $10^{-3}$ | 4.013 |
| Chlorite | 7.0 x $10^{-2}$ | 7.125 |
| Smectite | 2.3 x $10^{-3}$ | 68.240 |

The normalized adsorption capacity and storage capacity are calculated using:

$$\text{Normalized adsorption capacity of mineral x (\%)} = \frac{Adsorption\ capacity\ (x)}{\sum_x^n Adsorption\ capacities} * 100$$

$$\text{Storage capacity (\%)} = \frac{(Adsorption\ capacity\ of\ mineral\ x)*(mineral\ x\ wt.\%)}{\sum_x^n (Adsorption\ capacity\ of\ mineral\ *(mineral\ x\ wt.\%)} * 100$$

Table S2.2 Adsorption capacities, normalized adsorption capacities and storage capacities at 3 different DNA equilibrium concentrations ($C_{eq}$) based on adsorption isotherms measured in ASW

| | Equilibrium concentration: 0.0049 | | | Equilibrium concentration: 10 | | | Equilibrium concentration: 200 | | |
|---|---|---|---|---|---|---|---|---|---|
| | Adsorption capacity | Normalized Adsorption capacity | Storage capacity | Adsorption capacity | Normalized Adsorption capacity | Storage capacity | Adsorption capacity | Normalized Adsorption capacity | Storage capacity |
| | ug/mL | % | % | ug/mL | % | % | ug/mL | % | % |
| Quartz | 1.42E-05 | 0.014 | 0.901 | 0.027 | 0.202 | 9.943 | 0.255 | 0.226 | 9.062 |
| Feldspar | 1.81E-05 | 0.017 | 0.159 | 0.035 | 0.255 | 1.740 | 0.303 | 0.269 | 1.488 |
| K-feldspar | 9.07E-06 | 0.009 | 0.089 | 0.017 | 0.128 | 0.978 | 0.152 | 0.134 | 0.836 |
| Plagioclase | 9.07E-06 | 0.009 | 0.070 | 0.017 | 0.128 | 0.762 | 0.152 | 0.134 | 0.652 |
| amphibole | 2.96E-05 | 0.028 | 0.021 | 0.055 | 0.405 | 0.222 | 0.403 | 0.357 | 0.158 |
| pyroxene | 8.13E-05 | 0.078 | 0.297 | 0.135 | 0.998 | 2.844 | 0.596 | 0.528 | 1.222 |
| Illite | 4.15E-04 | 0.396 | 1.25 | 0.806 | 5.96 | 13.98 | 8.35 | 7.40 | 14.12 |
| biotite | 2.55E-02 | 24.33 | 9.47 | 2.30 | 17.03 | 4.93 | 9.73 | 8.63 | 2.03 |
| Chlorite | 7.33E-02 | 69.96 | 81.74 | 1.981 | 14.64 | 12.71 | 7.239 | 6.415 | 4.526 |
| smectite | 2.67E-03 | 2.549 | 4.05 | 4.061 | 30.01 | 35.47 | 72.36 | 64.12 | 61.57 |
| talc | 2.76E-03 | 2.632 | 2.11 | 4.127 | 30.50 | 18.15 | 13.61 | 12.06 | 5.83 |

Adsorption capacity for K-feldspar was set similar as to the value measured for feldspar. For biotite the average value for the adsorption capacity for muscovite, chlorite and talc was applied.

### Supporting info section 3. Distribution coefficients

To estimate the concentration of non-adsorbed DNA in seawater ($C_{eq}$) with various solid – water ratios we can use a distribution coefficient. Distribution coefficient is a ratio of DNA adsorbed to a specific mineral and the DNA concentration that is left in seawater after the equilibrium is reached:[1]

$$Distribution\ coefficient = \frac{Adsorption\ capacity}{c_{eq}}$$

Inserting the distribution coefficients into the equations for adsorption capacities of minerals gives us:

Distribution coefficient (Quartz) = $\frac{0.003}{1+0.006c_{eq}}$

Distribution coefficient (K-feldspar) = $\frac{0.002}{1+0.007c_{eq}}$

Distribution coefficient (Amphibole) = $\frac{0.006}{1+0.010c_{eq}}$

Distribution coefficient (Pyroxene) = $\frac{0.017}{1+0.023c_{eq}}$

Distribution coefficient (Talc) = $\frac{0.619}{1+0.040c_{eq}}$

Distribution coefficient (Illite) = $\frac{0.082}{1+0.005c_{eq}}$

Distribution coefficient (Kaolinite) = $\frac{0.301}{1+0.070c_{eq}}$

Distribution coefficient (Chlorite) = $0.73c_{eq}^{-0.57}$

Distribution coefficient (Smectite) = $0.40c_{eq}^{-0.03}$

Knowing the distribution coefficient we can calculate the fraction of DNA in seawater with different particle loadings.[1]

### Supporting info section 4. AFM

Strong or weak adsorption using AFM

In order to use AFM for imaging DNA molecules complexing with mica in liquid, cations with a strong ionic potential (charge/radius) such as Ni2+ are needed. Regardless of the immobilization facilitated by e.g. $NiCl_2$, drying the surface, has proven to provide information of the conformation of the DNA, inherent to the adsorbing liquid composition.[2] We added plasmid DNA in ASW to a mica surface and imaged the DNA in air to confirm adsorption. The plasmid are arranged in a range of conformations (collapsed, circular and coiled) (Fig. Sup1.a). Adding ASW solution to the surface in Fig. Sup1a, and scanning the surface in the liquid, we are unable to detect circular plasmids (Fig. Sup 1b). Instead, we observe several small and mobile spots which highlight that the DNA is weekly associated with the mineral surface and the interaction strength is not high enough to draw the DNA to the surface allowing it to be imaged with the AFM. Instead, the DNA is likely moved around by the AFM tip (as inferred by the striations observed in b and c). Reducing the salinity and the cation charge density of the solution to 1 mM $CaCl_2$ we are again unable to immobilize the DNA for imaging I the liquid and only observe mobile spots and striations (c). Despite decreasing the ionic strength of the solutions as well as the charge density of the cations available, drying the surface and scanning it in air reveal that the DNA did not desorb and confirm that the majority of the molecule was associated with the solution rather than with the surface while in liquid.

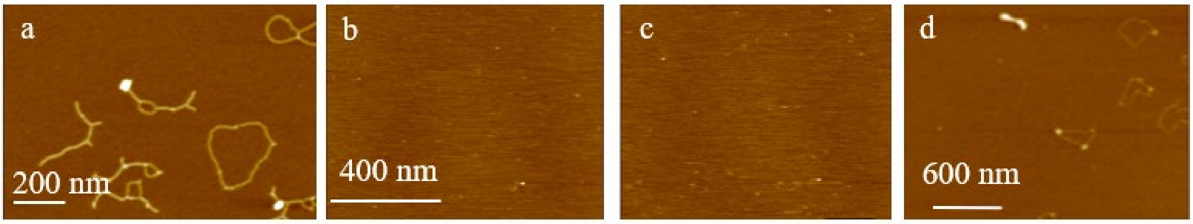


Figure S4.1. A) Plasmid DNA adsorbed in ASW and imaged in liquid. B). ASW added to the surface in A and imaged in the solution. C) ASW liquid was exchanged for 1 mM $CaCl_2$ and imaged in the liquid. D) The surface from C was dried and imaged in air.
